# Neural connectivity molecules best identify the heterogeneous clock and dopaminergic cell types in the *Drosophila* adult brain

**DOI:** 10.1101/2022.04.22.489196

**Authors:** Dingbang Ma, Nicholas Herndon, Jasmine Quynh Le, Katharine C. Abruzzi, Michael Rosbash

## Abstract

Our recent single cell sequencing of most adult *Drosophila* circadian neurons indicated striking gene expression heterogeneity, about 2-3 cells per clock neuron group. To extend this characterization to other adult fly brain neurons, we used the identical plate-based methods to generate single cell data from a similar subset of dopaminergic neurons. To minimize batch effects and to apply an additional sequencing strategy, we also assayed these two populations together with 10X Chromium. An unsupervised clustering algorithm indicates that dopaminergic neurons are comparably heterogeneous, suggesting that the transcriptomic diversity of adult fly brain neurons parallels its EM connectome. The results here further indicate that connectivity molecules like cell surface molecules best characterize all neuron groups. We suggest that these surprising features are general and make major contributions to neuronal identity and connectivity of the adult central brain as well as underlie the complex behavioral repertoire of *Drosophila*.

## Introduction

The fruit fly *Drosophila melanogaster* has a sophisticated repertoire of behaviors. Taken together with its remarkable genetic toolkit and tiny brain, it is an ideal model organism for studying the neuroscience of behavior. Somewhat at odds with this simplicity however is the extraordinary complexity of wiring and gene expression patterns of adult fly brain neurons, which is fundamental to understanding how these neurons form circuits and function to regulate behavior.

The fly brain wiring patterns have been illuminated by recent advances in genetics as well as light and electron microscopy (Scheffer et al., 2020), whereas advances in single cell RNA sequencing technologies have provided unprecedented insight into neuronal gene expression regulation (Svensson et al., 2018). These technologies have also been applied to many large groups of fly neurons, including optical lobes and whole-brain profiling (Brunet Avalos et al., 2019; Croset et al., 2018; Davie et al., 2018; Konstantinides et al., 2018; Kurmangaliyev et al., 2019; Li et al., 2022). These studies have led to the characterization of cellular plasticity during development as well as the de novo identification of cell type (Kurmangaliyev et al., 2020; Ozel et al., 2021), and they include a recent global collaboration effort, the Fly Cell Atlas project, to produce cellular gene expression maps of the entire fly (Li et al., 2022).

Single cell RNA sequencing after neuronal purification has also been applied to many smaller groups of fly brain neurons, including olfactory receptor neurons and olfactory projection neurons (Li et al., 2017; Li et al., 2020; McLaughlin et al., 2021; Xie et al., 2021). This approach has also revolutionized our view of the 150 adult brain circadian neurons, 75 in each hemisphere in the fly brain (Allada and Chung, 2010), within which a now well-defined transcription-translation feedback loop generates ∼24 hour periodicity (Top and Young, 2018).

These circadian neurons have historically been grouped based on anatomy. There are ventral lateral neurons, dorsal lateral neurons, and lateral posterior neurons. Four additional clock neuron groups were identified in the dorsal brain, including the DN1ps, DN1as, DN2s, and DN3s (Helfrich-Forster, 2004; Shafer et al., 2006). Single cell RNA sequencing of these clock neurons at different times of day revealed striking spatial and temporal gene regulation and substantially expanded the number of clock neuron groups, from about 8-10 to at least 17. Importantly, many of these high confidence clock neuron groups or clusters correspond to 2 or 3 neurons per hemisphere (Ma et al., 2021). It is however uncertain whether such transcriptomic heterogeneity is only true of these ca. 100 clock neurons or is characteristic of other fly central brain neuron groups. Such heterogeneity could even be common and contribute to the substantial central brain anatomical heterogeneity indicated by the hemibrain EM connectome.

To address this issue, we decided to characterize dopaminergic neurons (DANs). Immunohistochemistry of tyrosine hydroxylase (TH), an enzyme required for dopamine synthesis, had identified ∼282 neurons per adult fly brain. They were categorized into different anatomical clusters including four anterior groups (PAM, PAL, T1, and Sb) and 6 major posterior groups (PPL1, PPL2ab, PPL2c, PPM1, PPM2, PPM3) (Mao and Davis, 2009). A large body of evidence implicates DANs in the control of many different behaviors including locomotor activity, memory, arousal, aggression, and sleep (Alekseyenko et al., 2013; Andretic et al., 2005; Ganguly-Fitzgerald et al., 2006; Kume et al., 2005; Liu et al., 2012; Pendleton et al., 2002; Schwaerzel et al., 2003; Ueno et al., 2012). Consistent with their anatomical and functional diversity, DANs exhibit complex projection patterns: for example, 20 distinct DAN types each project axons to one or at most two mushroom body output neuron (MBON) compartments (Aso et al., 2014).

In this study, we generated around-the-clock single cell data from DANs by a modified CEL-seq2 method as we had previously done for clock neurons. To minimize batch effects and to apply an additional sequencing strategy, we labeled clock neurons and DANs in the same fly and assayed these two populations together with a droplet-based method (10X Chromium). An unsupervised clustering algorithm identified 43 high confidence clusters in the pooled dataset, and all of the previously identified clock neuron clusters were matched unambiguously in the current dataset. Using these clock neuron clusters as a benchmark, we show that dopaminergic neurons are comparably heterogeneous and that cell surface molecules and neuronal connectivity molecules more generally are prominent features of adult brain neuron identity, like during development of the visual system and projection neurons (Kurmangaliyev et al., 2020; Li et al., 2017; Ozel et al., 2021; Xie et al., 2021).These data and others suggest that these features may be general properties of neurons within the *Drosophila* central brain.

## Results

### Single-cell sequencing of clock and dopaminergic neurons by plate and droplet methods

To further characterize the gene expression heterogeneity in *Drosophila* neurons, we performed single cell RNA sequencing of DANs labeled by TH-GAL4 (Mao and Davis, 2009), which are comparable in number to clock neurons labeled by Clk856-GAL4 (Gummadova et al., 2009). As clock gene expression is very low or absent in DANs (Abruzzi et al., 2017), they should be representative of at least some non-circadian fly brain neurons.

We first characterized DANs with the same modified CEL-Seq2 method we used with clock neurons (Hashimshony et al., 2016; Ma et al., 2021) and even assayed different times of day by collecting flies around the clock (Figure 1A). Although these data could be compared with previous clock neuron data (Ma et al., 2021), we added an additional approach: TH-GAL4 and Clk856-GAL4 were combined into a stable line (Supplementary Figure 1A) to profile clock neurons and DANs together at two LD (Light:Dark) time points, ZT02 and ZT14; this stable line avoids batch effects. We also changed the assay method: the single cell RNA sequencing of this combined line was done with 10X Chromium chemistry. This provided another point of comparison with the clock neurons, modified CEL-Seq2 vs 10X sequencing.

**Figure 1.**
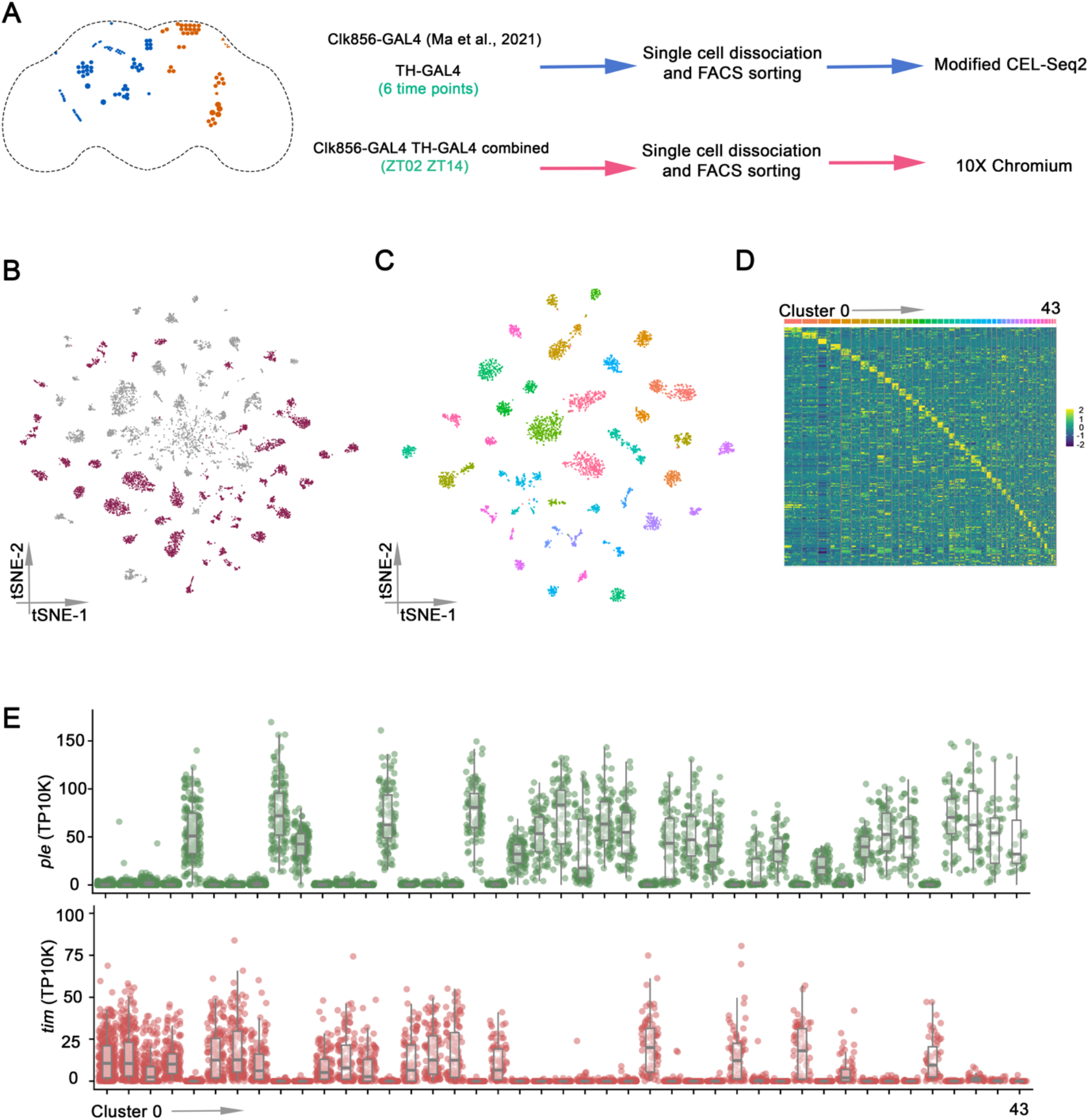
Single cell RNA sequencing of *Drosophila* clock neurons and dopaminergic neurons by plate and droplet-based methods. (A) Schematic workflow of single cell RNA sequencing data generation. The clock neurons (orange) and DANs (blue) are shown in the schematic depiction on the left panel. The data from clock neurons in Light:Dark conditions were from our previous publication. EGFP-labeled dopaminergic neurons were sorted into 384-well plates by fluorescence-activated cell sorting (FACS) and single cell RNA sequencing libraries were prepared by modified CEL-seq2 method. For droplet-based method, TH-GAL4 and Clk856-GAL4 were combined to a stable line in order to assay dopaminergic neurons and clock neurons simultaneously, fly brains were dissected and dissociated prior to droplet encapsulation of individual cells with barcoded beads in 10X Chromium. (B) t-Distributed stochastic neighbor embedding (t-SNE) plot showing the 9025 cells grouped into 70 clusters. High confidence clusters are shown in purple. (C) t-SNE plot of 4543 *Drosophila* clock and dopaminergic neurons in 43 high confidence clock and dopaminergic neuron clusters from Light: Dark conditions. The clusters are colored by their cell types. (D) Heatmap showing the expression levels of the top 5 differentially expressed genes (rows) in cells (columns). Clusters are ordered by size and are represented by different colors on top of the heatmap. (E) Dot plot showing *tim* (bottom) and *ple* (top) expression in all clusters. Gene expression levels for each cell were normalized by total expression level, we report transcripts per 10 thousand transcripts (TP10K). Clusters are ordered by size.

To catalog and compare single-cell gene expression in clock neurons and DANs, we combined the single cells from the plate- and droplet-based methods in the current study with the previous CEL-Seq2 clock neuron LD data (Ma et al., 2021) (Supplementary Figure 1B). 9025 cells remained after stringent filtering (Supplementary Figure 1C). Next, single cells were clustered by an unsupervised clustering method that produced 70 distinct clusters (Figure 1B). Although there was some variability, cells from both CEL-Seq2 and 10X were present in all clusters (Supplementary Figure 2A), and similar numbers of genes were identified in most clusters (Supplementary Figure 2B).

To identify clusters most likely to correspond to specific clock neurons and DANs, we filtered them to ensure that each cluster has cells from all timepoints and from both experimental methods (Supplementary Figure 2C-D) and that the clusters are minimally contaminated with poor-quality cells (see Methods). This resulted in 43 high-confidence clusters (Figure 1C), and each cluster showed enriched maker gene expression (Supplementary Figure 2E). Differential gene expression revealed that every cluster has a unique marker gene expression profile, supporting the validity of the strategy (Figure 1D).

To distinguish between clock and DAN clusters, we characterized the expression of *timeless* (*tim*) and *pale* (*ple*), which are hallmark genes for these two neuron groups. Consistent with previous results showing that clock gene expression is very low in DANs (Abruzzi et al., 2017), we found that *tim* and *ple* expression is mutually exclusive and defines 19 clock and 24 DAN clusters (Figure 1E). Other DAN-positive genes are co-expressed with *ple* and other clock genes are co-expressed with *tim* (Supplementary Figure 3). *tim* mRNA is also cycling with its characteristic gene expression peak at ZT14-ZT18 in all clock clusters (Supplementary Figure 4). Moreover, genome wide cycling gene expression analysis identified many cell-type specific cycling transcripts in clock clusters but only a minimal number of oscillating transcripts in DAN clusters (data not shown), consistent with previous results (Abruzzi et al., 2017).

### Identifying clock neuron and DAN clusters

Our previous study identified at least 17 high confidence clock neuron groups with striking spatial and temporal regulation of gene expression (Ma et al., 2021). To correlate these groups with our new dataset, we first characterized the expression of known clock neuron marker genes. For example, the neuropeptide genes *Pigment-dispersing factor* (*Pdf*), *Trissin* and *CCHa1* were expressed in 3 different clock neuron groups: 8 LNv, 2 LNd and 2 DN1a clock neurons, respectively (Fujiwara et al., 2018; Helfrich-Forster, 1995; Ma et al., 2021). Similarly, each of these neuropeptide transcripts corresponds to a single clock neuron cluster in the new single cell RNA sequencing data (Figure 2A). More generally, this marker gene strategy along with core clock gene expression could match almost all new clock neuron clusters to the previous assignments (Figure 2B). The only exceptions are the new cluster 2, which contains cells from multiple clock neurons groups (Supplementary Figure 5A), and the new cluster 25; the latter is an extra DN1p cluster, reflecting 2 DN1p clusters where there was previously only one.

**Figure 2.**
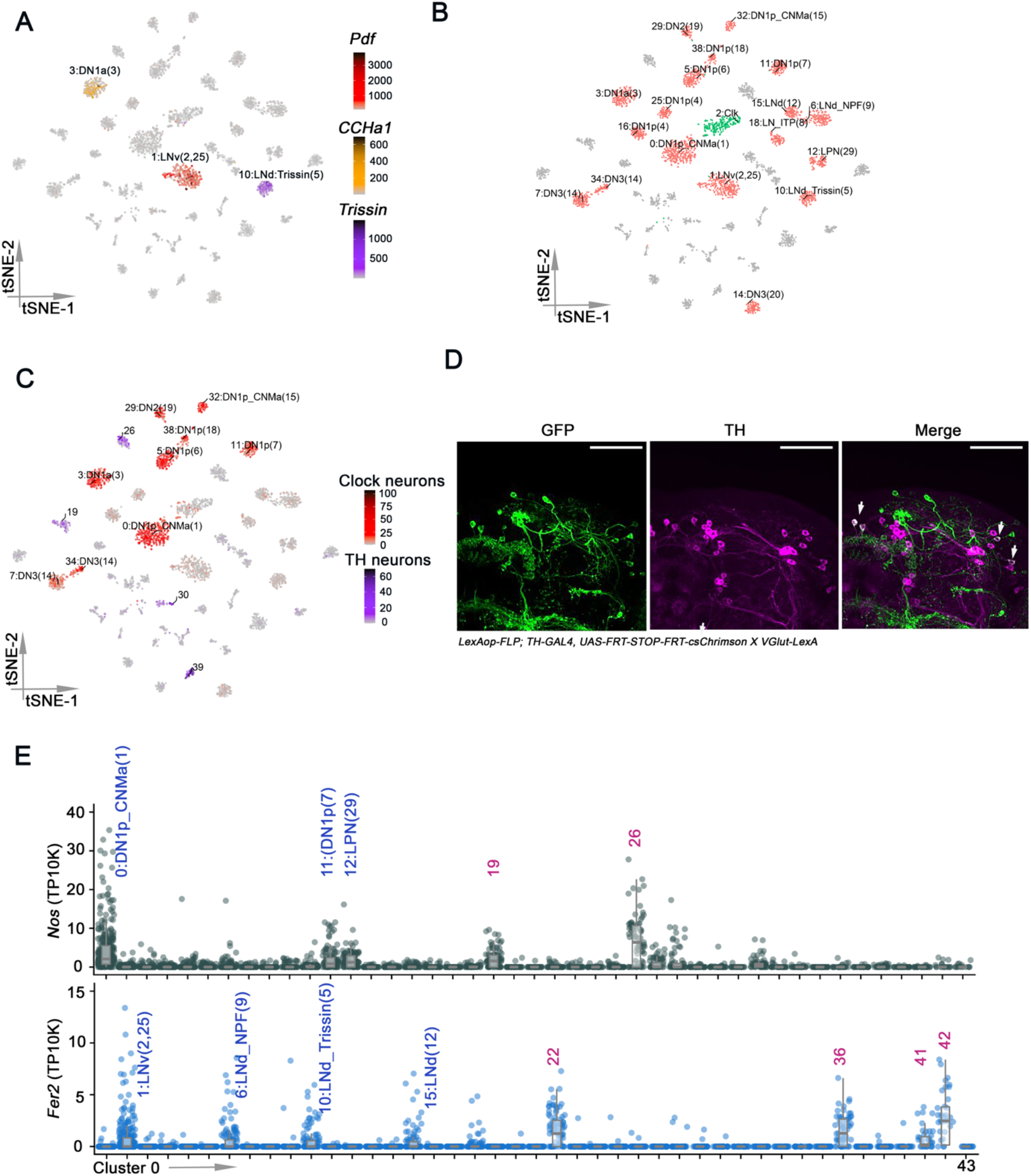
Identifying clock neurons and dopaminergic neurons clusters. (A) t-SNE plots showing *Pdf* (red)*, Trissin* (purple) *and CCHa1* (yellow) expression in all clusters. Each cell is colored by the expression (color bars, TP10K) with gray indicating low expression and black indicates the highest expression. (B) t-SNE visualization of the annotated clock clusters. Each clock neuron cluster retains its original identifying number in the parentheses as it was previously reported. The clusters in red were all previously identified and cluster 2 (green) is new in the current study. (C) t-SNE plots showing *VGlut* expression in all clusters. Each cell is colored by the expression of *VGlut* (color bars, red and purple represents *VGlut* expression in clock neurons and DANs, respectively, TP10K), with gray indicating low expression and black indicating the highest expression. (D) Confocal stack of images showing antibody staining for GFP (left), TH (middle) and the merged image (right) in *VGlut-LexA* > *LexAop-FLP; TH-GAL4* > *UAS-FRT-STOP-FRT-Cschrimson.venus* fly brains. Two PPM1 and two PPL1 neurons are both GFP- and TH-positive. Scale bars represent 50 μm. (E) Dot plot showing *Fer2* (bottom) and *Nos* (top) expression in all clusters. Gene expression levels for each cell were normalized by total expression level, we report transcripts per 10 thousand transcripts (TP10K). Clusters are ordered by size.

An interesting marker is the vesicular glutamate transporter *VGlut* mRNA, which identifies glutamatergic neurons. It is expressed in 9 dorsal clock neuron clusters (Figure 2C, red), which corresponds precisely to the prior number of glutamatergic clock neuron groups (Hamasaka et al., 2007; Ma et al., 2021). *VGlut* is also expressed in 4 dopaminergic clusters (Figure 2C, purple), consistent with previous findings that glutamate and dopamine are co-expressed in some larval and adult fly brain neurons (Aguilar et al., 2017; Brunet Avalos et al., 2019). In contrast to flies, glutamate is the major excitatory neurotransmitter in mammals and is also co-expressed with dopamine in some mammalian neurons (Chuhma et al., 2004; Hnasko et al., 2010; Hnasko and Edwards, 2012).

To further verify the expression of *VGlut* in fly DANs we used a method that allows genetic access to the neurons at the intersection of a LexA and GAL4 line. We expressed UAS>stop>CsChrimson.venus, a FLP-dependent conditional reporter, in TH-GAL4 in addition to LexAop-FLP in a VGlut-LexA knock-in line. Cells that express both the conditional reporter and FLP were then labeled with Cschrimson.venus. CsChrimson.venus expression was restricted to only a few DANs, which include the two PPM1 and two PPL1 neurons (Figure 2D); they were the strongest and most reproducible double-positive cells and had not been previously identified as glutamatergic.

To further identify DAN clusters, we examined the expression of several known dopaminergic marker genes. Nitric oxide is a cotransmitter in a subset of dopaminergic neurons and acts antagonistically to dopamine (Aso et al., 2019). Indeed, bulk RNA sequencing and immunohistochemistry showed that *Nos* (nitric oxide synthase) is expressed in PPL1-γ1pedc and PAM-γ5 neurons (Aso et al., 2019). The data here indicate that *Nos* is expressed in two DAN clusters (Cluster 19 and 26), which suggests that they correspond to PPL1-γ1pedc and PAM-γ5 neurons. This further suggests that the two other glutamate-expressing DAN clusters, 30 and 39, correspond to PPM1 and PPL1 (Figure 2D). *Nos* mRNA has not been described in clock neurons, but the gene is apparently comparably expressed in 3 DN1p clock neuron clusters, cluster 0, 11 and 12 (Figure 2E, top).

*48-related-2* (*Fer2*) encodes a transcription factor, which is expressed in DANs in the protocerebral anterior medial (PAM) cluster and required for their development and survival (Bou Dib et al., 2014). Although immunohistochemistry of tyrosine hydroxylase (TH) indicates that there are about 100 PAM DANs, the *TH-GAL4* driver only identified 13 PAM cells per hemisphere (Mao and Davis, 2009). As there are 4 DAN clusters with enriched *Fer2* expression (Figure 2E, bottom), which presumably contain these 13 PAM DANs, suggesting an average of about 3 PAM DANs per cluster. There are apparently 4 clock clusters that also express *Fer2* (Figure 2E, bottom).

These marker genes suggest assignments of 8 DAN clusters to known dopaminergic neurons. To assign additional clusters, we compared cluster gene expression with previously published transcriptomes from several different DANs subgroups (Aso et al., 2019). Most of these comparisons were unsuccessful, perhaps because most subgroups were PAMs and they are very underrepresented by TH-GAL4. However, an otherwise uncharacterized DAN cluster, #28, is highly correlated with PPL1-α3 and PPL1-γ2 α’2 neurons (Supplementary Figure 5B). This assigns a 9th cluster and hints that this single cluster may not be homogeneous, i.e., that the dopaminergic neurons are even more heterogeneous than indicated by this analysis.

### Neuropeptide expression in clock neurons and DANs

To further characterize the clock neurons and DANs, we assayed differential gene expression using a negative binomial generalized linear model. Consistent with previous findings, Gene Ontology (GO) analysis indicates that clock neuron transcriptomes are enriched for the term neuropeptide hormone activity (Abruzzi et al., 2017; Ma et al., 2021); indeed, it was the top term (Figure 3A), and many clock neurons express a complex combination of neuropeptides (Figure 3B). This same term is present in DANs but lower down on its list (Figure 3A), and only a handful of specific neuropeptides were identified in DANs compared to clock neurons (Figure 3B).

**Figure 3.**
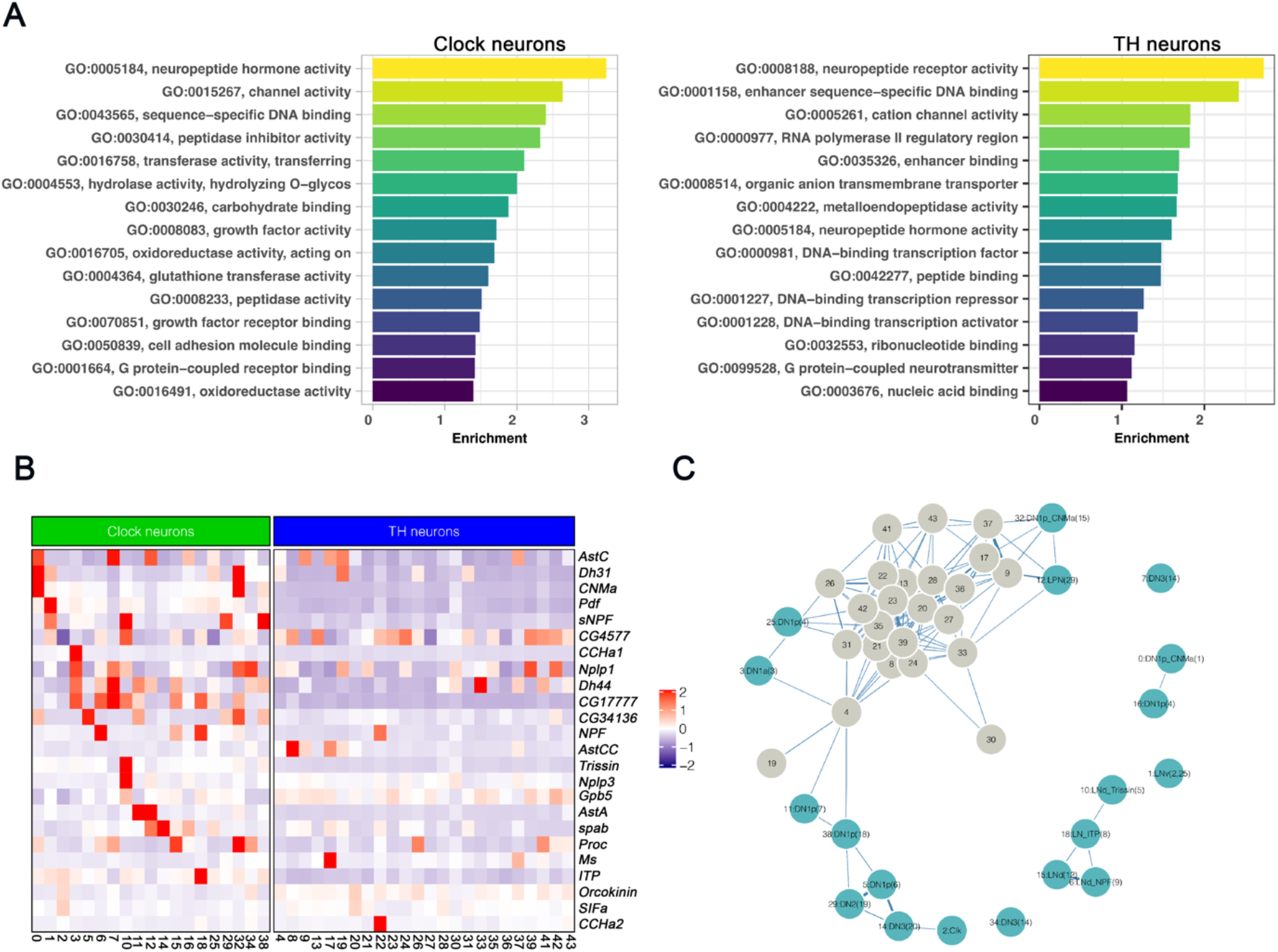
Neuropeptides expression in clock neurons and dopaminergic neurons clusters. (A) Gene ontology (GO) analysis of enriched marker genes found in the clock neuron and DAN clusters. The enriched GO terms were ranked by their relative enrichment. (B) Heatmap showing the expression levels of neuropeptides in clock neurons (left panel) and DANs (right panel). Red indicates high expression and purple indicates low expression. (C) Gene expression correlation of neuropeptides in clock neurons and DANs. We calculated the Spearman’s correlation coefficients between expression patterns of neuropeptides across different clock neuron and DANs cell types, the result is visualized in a force embedded layout. Each cluster is represented by a node with the width of the edge representing the strength of the gene expression.

Nonetheless, 4 neuropeptides -- *Dh44*, *Nplp1*, *Glycoprotein hormone beta 5* (*Gpb5*) and *Proctolin* (*proc*) -- have been reported to be expressed in up to 21% of DANs (Crosest et al., 2018), but our data indicate that only *Dh44* mRNA is clearly expressed in the TH-Gal4 DANs (Supplementary Figure 6A). *Dh31* mRNA is also enriched in one DAN cluster. As mentioned above, it probably corresponds to PPL1-γ1pedc or PAM-γ5 (Supplementary Figure 6B). *Dh31* is also expressed in the two previously described clock neuron clusters (Kunst et al., 2014; Ma et al., 2021) (Supplementary Figure 6B); The transcript encoding the neuropeptide *Ms* is similarly detected in a single DAN cluster, #17 (Supplementary Figure 6C).

Several dopaminergic neurons do however express more than one neuropeptide mRNA: *AstC* is expressed along with *Ms* in cluster 17 (Supplementary Figure 6C-D); *Dh31* and *AstC* are co expressed in cluster 19 (Supplementary Figure 6B, D). Surprisingly, this likely PPL1-γ1pedc or PAM-γ5 cluster co-expresses glutamate and *Nos* as well as dopamine. Despite these exceptions, neuropeptide transcript expression in the DANs appears limited and may be influenced by incomplete coverage by TH-Gal4 (see Discussion).

To further compare neuropeptide expression between clusters and between clock neurons and DANs, we calculated the Spearman gene expression correlation based on the average expression of neuropeptides within each cluster. The results were then visualized with a forced embedded layout (Figure 3C). The separation of the individual clock neuron clusters indicates that differential neuropeptide expression defines very successfully these clusters but much less well the poorly separated DAN clusters. This conclusion may simply reflect the many more neuropeptides in clock neurons than in DANs (Figure 3B).

### Neural connectivity molecules best identify DANs as well as clock neurons

To identify other genes that might underlie cell type definition of DANs as well as clock neurons, we turned to the highly variable genes that were used during our data integration and clustering analysis. We computed 3000 highly variable genes based on the different raw gene expression datasets separated by time points and methods, of which 338 were expressed and variable in all datasets and then used for further downstream analysis (Figure 4A). Hierarchical clustering analysis based on these 338 highly variable genes grouped clock neurons and DANs into different clusters (Supplementary Figure 7A). We next cataloged these highly variable genes into different functional groups (Figure 4B): the top 3 are transcription factors (TFs), cell surface molecules (CSMs) and G protein-coupled receptor (GPCRs).

**Figure 4.**
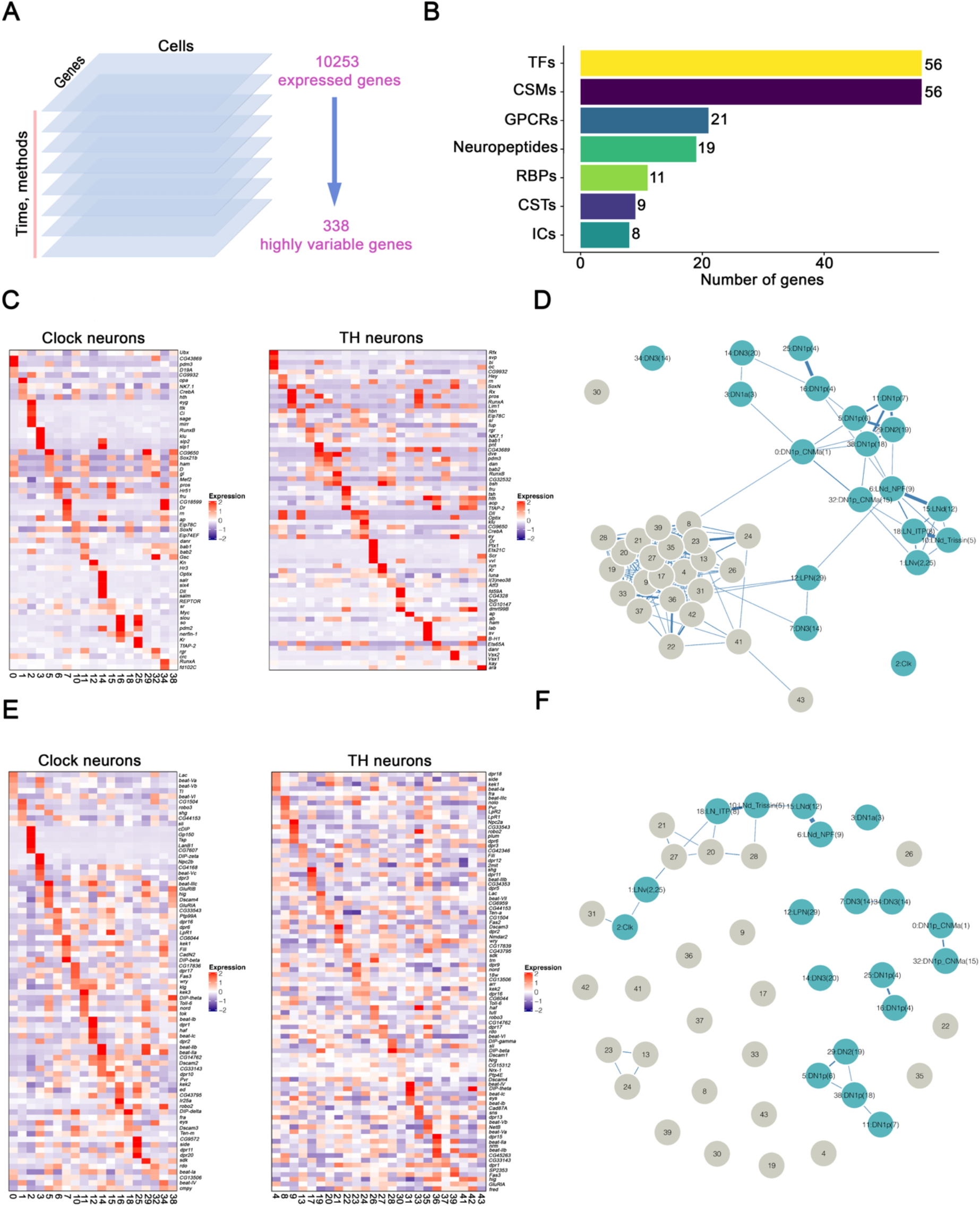
Transcription factors and cell surface molecules expression in clock neurons and dopaminergic neurons. (A) Schematic workflow of data integration and clustering analysis. We first separated the single cell gene expression results by time points and methods. In each dataset, 3000 highly variable genes were calculated and only the conserved variable genes (338 genes) were used for final single cell clustering. (B) Bar plot showing the number of highly variable genes from different gene groups including transcription factors (TFs), cell surface molecules (CSMs), G protein coupled receptors (GPCRs), neuropeptides, RNA binding proteins (RBPs), chemical synaptic transmission related genes (CSTs) and ion channels (ICs). (C, E) Heatmaps showing the expression levels of transcription factors (C) and cell surface molecules (E) in clock neurons and DANs. (D, F) Gene expression correlation of transcription factors (D) and cell surface molecules (F) in clock neurons and DANs. We calculated the Spearman’s correlation coefficients between expression patterns of neuropeptides across different clock neuron and DANs cell types; the result is visualized in a force embedded layout. Each cluster is represented by a node with the width of the edge representing the strength of the gene expression.

Spatial regulation of TF expression is prominent in clock neurons and may contribute to the robust temporal oscillation of many transcripts (Ma et al., 2021). Each DAN cluster also expresses a specific combination of TF genes, very similar to the observed patterns in clock neurons (Figure 4C). However, the gene expression correlation analysis indicates that TFs do not separate the DANs as well as the clock neurons (Figure 4D), and the clock neurons appear somewhat less well defined by TFs than by neuropeptides: the clock neuron clusters are closer together and have more links in Figure 4D than in Figure 3C.

The highly variable genes contain an almost identical number of CSMs as TFs (Figure 4B). Although CSMs are critical in mediating interactions between cells in the developing nervous system (Zinn and Ozkan, 2017), these molecules do not have a well-defined general role in the adult nervous system. Yet the CSM heatmap indicates impressive cluster-specific expression, in DANs as well as in clock neurons (Figure 4E). Moreover, the gene expression correlation analysis indicates that all single cell clusters in both populations are very well separated by CSM expression (Figure 4F). Notably, this class of molecules includes the Dpr (Defective proboscis extension response) and DIP (Dpr-interacting protein) protein families. In agreement with previous findings on clock neurons (Ma et al., 2021), each clock neuron and DAN cluster expresses a specific combination of Dprs (Supplementary Figure 7B), and this is also the case for DIPs (data not shown).

GPCRs are another category of neuron connectivity molecules expressed in the highly variable genes (Figure 4B). GPCRs are 7-transmembrane receptors that interact with many different stimuli including neurotransmitters and neuropeptides and play an important role in the physiology and function of neurons (Brody and Cravchik, 2000). More than 2/3 of the 124 GPCR genes encoded by the *Drosophila* genome are expressed in adult clock neurons, and these molecules alone can define clock neuron cell type (Schlichting et al., 2021). The data here indicate that these transcripts define equally well the DAN clusters (Figure 5A-B).

**Figure 5.**
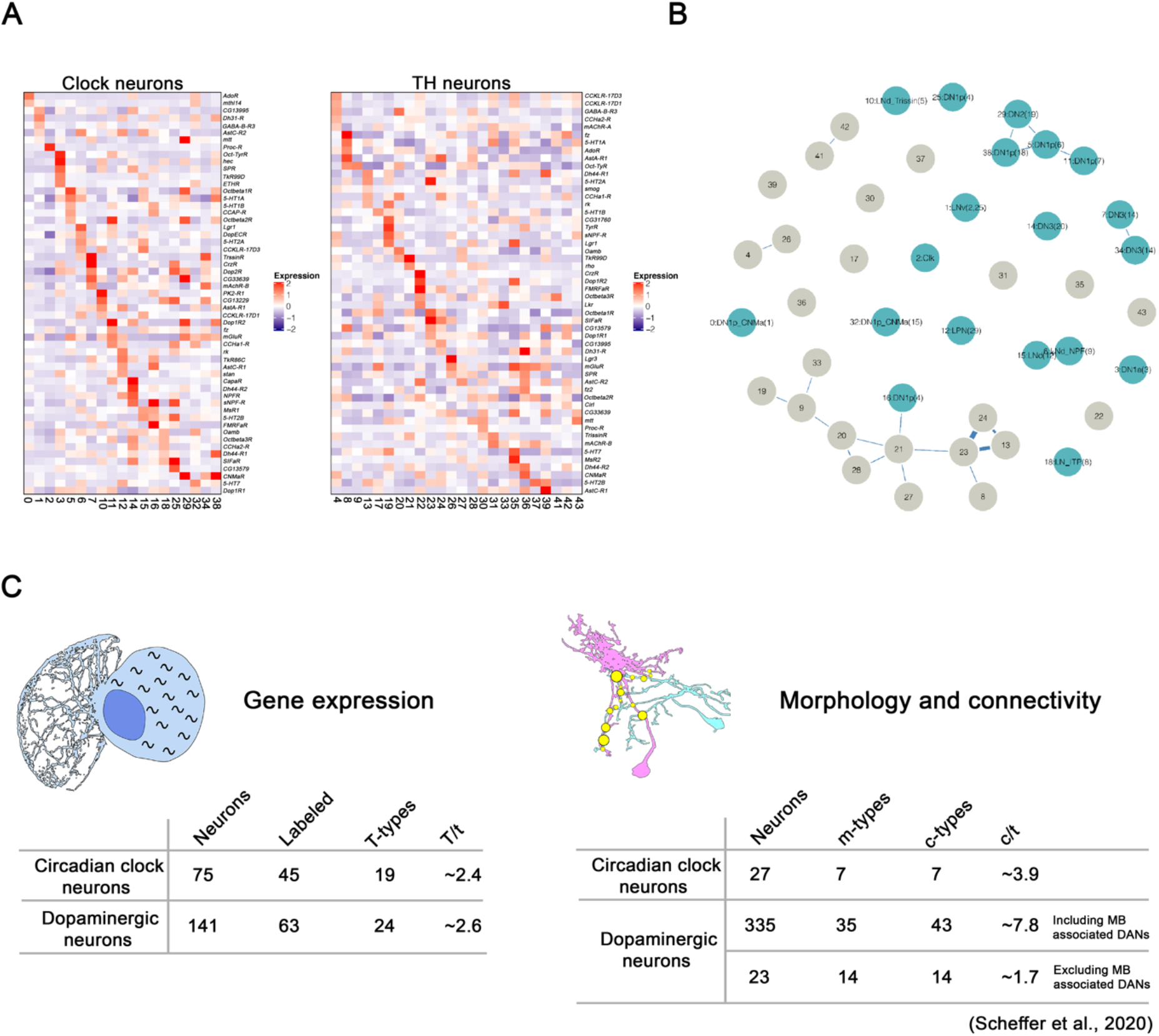
GPCR expression in clock neurons and dopaminergic neurons. (A) Heatmaps showing the expression levels of GPCRs in clock neurons and DANs. (B) Gene expression correlation of GPCRs in clock neurons and DANs. We calculated the Spearman’s correlation coefficients between expression patterns of neuropeptides across different clock neuron and DANs cell types; the result is visualized in a force embedded layout. Each cluster is represented by a node with edge width representing the strength of the gene expression. (C) A summary of identified gene expression clusters (T/t, average number of cells per transcriptomic type) in the current study and connectivity types (c/t, average number of cells per connectivity type) from the hemi-brain EM dataset, m-types are the number of morphology types. The cell number includes some neurons on the contralateral side, they represent the number of cells that are included in the clustering, but not the number of neurons per brain side (Scheffer et al., 2020).

In summary, the data here indicate that neuron connectivity molecules -- neuropeptides (for clock neurons), CSMs and GPCRs -- constitute an important class of neuron identification molecules in the adult fly nervous system, at least for clock neurons and DANs.

## Discussion

There are about 75 *Drosophila* clock neurons on each side of the adult fly brain. They are defined by their common features, most notably cycling circadian gene expression. They had been divided into small subgroups based on their distinctive anatomical locations and projection patterns, and some had also been distinguished based on immunohistochemistry, namely the expression of specific neuropeptides, as well as function. However, the advent of single cell RNA sequencing and the ability to comprehensively profile gene expression have been game changers for neuron characterization (Li, 2021; Mohr et al., 2021); indeed, our profiling of fly clock neurons with CEL-seq2 dramatically increased the number of clock neuron subgroups compared to previous definitions. Other features were also notable, for example the spatial and temporal regulation of cell-surface proteins (Ma et al., 2021).

To bring some perspective to our previous clock neuron characterization, we assayed here DANs. They were chosen only because they are about as numerous as clock neurons in the adult fly brain and because these neurotransmitter-expressing neurons might be very different. The two populations are however quite similar.

We first characterized DANs with CEL-seq2, identically to our previous clock neuron work. We then profiled together the transcriptomes of 1979 DANs and 2564 clock neurons (4543 high quality single cells in total), purified from a single strain expressing GFP in DANs as well as in clock neurons. By assaying the two populations together, batch effects were eliminated; we also used a different sequencing method for this dual assay, 10X Chromium. This strategy allowed our previous characterization of clock neurons with CEL-seq2 to serve as a benchmark against which the DAN data, the new clock neuron data and the 10X sequencing approach could be compared. Indeed, the 10X sequencing performed very well compared to CEL-seq2. Expressed genes per cell were almost identical, and CEL-seq2 had only about twice as many transcripts/gene (data not shown).

It is reassuring that an unsupervised clustering algorithm identified all previously defined clock neuron groups in the pooled data set. There are 19 clock neurons clusters in the new data, and we had previously identified 17 (Ma et al., 2021). As the Clk856-GAL4 driver used for clock gene expression expresses in 45 neurons/hemisphere, 45/17-19 clusters is ∼2.5 neurons/cluster. Using this complexity as a benchmark, the results show that dopaminergic neuron heterogeneity is nearly identical. The TH-GAL4 dopaminergic driver expresses in 63 dopaminergic neurons per hemisphere, which were categorized into 24 TH clusters with an average of 63/24 ∼2.6 neurons per cluster. If this is characteristic of all 140 dopaminergic neurons per hemisphere, there are at least 50-60 different DAN transcriptome types per hemisphere. This level of heterogeneity echoes the classification of neuron morphology and connectivity based on the hemi-brain EM dataset (Scheffer et al., 2020) (Figure 5C). In this context, Waddell and colleagues have identified 20 PAM-γ5 subtypes by anatomy (Otto et al., 2020), and it would not be surprising if many of these anatomical subtypes are also discrete by transcriptional profiling. Consistent with this notion are recent anatomy and connectome analyses of the 6 LNd clock neurons: there are 4 different groups, which precisely match our previous classification based on gene expression (Ma et al., 2021; Shafer et al., 2022; Yao et al., 2016).

This complexity of the adult fly brain transcriptomes is not predicted by previous results. Although there is even greater diversity in *Drosophila* visual and olfactory neurons during developmental wiring, it is not maintained in the adult, especially between visual system neurons that only differ in their connectivity and perform very similar functions (Kurmangaliyev et al., 2019; Kurmangaliyev et al., 2020; Li et al., 2017; Li et al., 2020; Ozel et al., 2021). These systems may therefore not be characteristic of most central brain neurons. Moreover, whole fly brain single cell sequencing efforts identified only a single DAN and a single clock neuron cluster (Supplementary Figure 8A-C). The very recent whole fly cell atlas indicates more heterogeneity, but the circadian cells are predominantly photoreceptor, cone and epithelial cells (Supplementary Figure 8D, E). (Croset et al., 2018; Davie et al., 2018; Li et al., 2022). It is likely that these data are too shallow and specific cells insufficiently numerous to reveal the many clock and dopaminergic central brain neuron groups. Transcriptomic definition of cell type may require at least 1 million cells just from the adult fly central brain, many more than what has been done to date in whole brain and whole fly studies (Brunet Avalos et al., 2019; Croset et al., 2018; Davie et al., 2018; Li et al., 2022). This would be sufficient to profile 30-100 copies of the perhaps 10,000-30,000 different cell types indicated by the EM connectome of the fly brain. An alternative approach is to use cell or nuclear purification from a very large number of specific drivers as we have done here for two drivers.

Both drivers, Clk856-GAL4 and TH-GAL4, fail to express in a substantial fraction of their expected patterns, presumably because they are simply missing important regulatory information that normally dictates *clock* or *pale* (TH=tyrosine hydroxylase) expression. These missing neurons may account for some of the data shortcomings as well as the few differences between the clock neuron and DAN populations. The Clk856-GAL4 driver expresses in only a few DN3 clock neurons despite expressing in all clock neurons with well-characterized functions. The enigmatic DN3s constitute about half of the clock neuron population, 30-40 cells per hemisphere. Similarly, the TH-Gal4 driver fails to express in more than half of the fly brain DANs, about 78/141 neurons per hemisphere. These missing DANs include most of the numerous protocerebral anterior medial (PAM) DANs, which are very underrepresented in TH-GAL4; it expresses in only 13 PAMs per hemisphere rather than the known 100 PAMs per hemisphere. It is possible that these missing neurons can explain some of the differences between the two populations. For example, there is much more neuropeptide transcript expression in clock neurons than in DANs (Figure 3). This is presumably why neuropeptide expression can separate clock neuron clusters much more successfully than DAN clusters (Figure 3C). This distinction may be less striking if DN3 clock neurons express relatively few neuropeptide transcripts compared to Clk856-GAL4 positive clock neurons and/or if the missing DANs express many neuropeptide transcripts compared to the DANs identified by TH-GAL4. In any case, it is uncertain whether the current characterization of neuropeptide expression in clock neurons or in DANs reflects the general case for the fly brain if indeed there is one. It is also uncertain whether the mammalian SCN is like fly brain clock neurons a richer source of neuropeptide expression than elsewhere in the mammalian brain (Xu et al., 2021).

Transcription factors separate clock neuron clusters more successfully than DAN clusters (Figure 4D). In contrast to neuropeptide transcript expression however, there is no indication that transcription factor expression is different between DANs and clock neurons; the heatmaps appear similar, and the relevant GO terms are even more highly ranked in DANs than in clock neurons. The gene expression correlation plot may then reflect that fact that transcription factor expression is more discrete in clock neurons than in DANs, for example that transcription factor specification may be more combinatorial in DANs than in clock neurons. This level of detail may not appear in the heatmaps, which are based on the top 5 transcription factors.

We were surprised to discover that cell surface protein transcripts, including GPCRs and CSMs, are the best definers of neuron identify, for DANs as well as for clock neurons. We had previously found that GPCRs can identify clock neurons (Schlichting et al., 2021), and these data extend this conclusion to many other classes of cell surface proteins as well as to DANs. CSMs are critical for nervous system wiring during development, best illustrated perhaps by their remarkably complex expression patterns during fly visual system development. As an example, the Dpr proteins have affinity for specific partner DIP proteins, which together help drive synapse specificity during visual system development (Sanes and Zipursky, 2020; Zinn and Ozkan, 2017).

The important role of connectivity molecule expression in neuron definition is especially surprising in the case of neuropeptides and GPCRs. This is because there are relatively few of these genes in the set of 338 highly variable genes that are used to cluster the cells (Figure 4B). Neuropeptides do very well only for clock neurons, reflecting no doubt the relative abundance of these genes in this cell type, whereas GPCRs do equally well in defining DANs as well as clock neurons. The success of GPCRs recalls the area code hypothesis, a transmembrane receptor cell surface code for embryo assembly (Dreyer, 1998), as well as the success of neuropeptides and GPCR pairs in cortical neuron definition (Smith et al., 2019). Notably, GPCRs appear to define neuron type about as well as the much larger number of CSMs. CSMs then are superior to transcription factors, indicating that the discrete expression of CSMs, GPCRs and neuropeptides (in the case of clock neurons) is a major feature of neuron-specific gene expression.

These data further suggest that these molecules are important to maintain wiring specificity and/or synaptic strength in the adult brain as well as create it during development. Their expression may also change with experience, environment or time, which suggests that aspects of neuronal cell surface expression as well as anatomy of much of the adult brain are plastic (Fernandez et al., 2008; Song et al., 2021). This is perhaps best illustrated by the now classic circadian regulation of circadian neuron projection pattern (Fernandez et al., 2008) as well as the more recently described temporal regulation of cell surface molecule expression in clock neurons (Ma et al., 2021). There are also circadian behavioral changes dependent on specific GPCR expression in specific clock neurons (Schlichting et al., 2021). Although these observations need to be extended to morphology as well as to DANs and eventually to other classes of adult neurons, the findings here emphasize the importance and broad reach of neuron connectivity molecules, which extend from anatomy and brain wiring to their specific gene expression patterns within individual adult brain neurons.

## Key resources table

| Reagent types | Designation | Source or reference | Identifiers | Additional information |
| --- | --- | --- | --- | --- |
| Genetic reagent<br>(*D. melanogaster*) | UAS-Stinger | BDSC | RRID:BDSC_84277 |  |
| Genetic reagent<br>(*D. melanogaster*) | Clk856-GAL4 | (Gummadova et al. 2009) | Flybase: FBtp0069616 |  |
| Genetic reagent<br>(*D. melanogaster*) | UAS-EGFP | BDSC | RRID:BDSC_5428 |  |
| Genetic reagent<br>(*D. melanogaster*) | UAS-FRT-STOP-FRT-CsChrimson.mVenus and LexAop-FLP | (Guo et al. 2018) | RRID:BDSC_84277 |  |
| Genetic reagent<br>(*D. melanogaster*) | VGlut-LexA | (Deng et al. 2019) |  |  |
| Genetic reagent<br>(*D. melanogaster*) | TH-GAL4 | (Abruzzi et al. 2017) |  |  |
| Antibody | Anti-TH<br>Rat monoclonal | IMMUNOSTAR |  | 1:1000 |
| Antibody | anti-GFP<br>Chicken | Abcam | RRID:<br>AB_300798 | 1:1000 |
| Antibody | Goat anti-mouse<br>polyclonal | ThermoFisher | RRID:<br>AB_2536185 | 1:200 |
| Antibody | Goat anti-Chicken<br>polyclonal | ThermoFisher | AB_2534096 | 1:200 |
| Software,<br>algorithm | FIJI | https://fiji.sc/ |  |  |
| Software,<br>algorithm | RStudio | https://rstudio.co<br>m | RRID:<br>SCR_000432 | Version 1.2.5033 |
| Software,<br>algorithm | Custom code | This paper |  | GitHub |

## Methods

### Fly stains and rearing

Flies were reared on standard cornmeal medium with yeast under 12:12 Light: Dark conditions at room temperature. The fly lines used in this study are listed in the key resource table. Equal numbers of males and females were used in all the single cell RNA sequencing library preparation.

### Fluorescence activated cell sorting

We used EGFP to label the targeted neurons. Flies were entrained in 12:12 Light: Dark cycles and 25 °C conditions for 3 days prior to dissection. For CEL-Seq2 experiments, time points were taken every 4 hours within a day, and ZT02 (2 hours after lights on) and ZT14 were used for the 10X Chromium experiment. Fly brains were dissected in cold dissection saline (HEPES-KOH 9.9 mM pH7.4, NaCl 137 mM, KCl 5.4 mM, NaH2PO40.17 mM, KH2PO40.22 mM, glucose 3.3 mM and sucrose 43.8 mM) with neuronal activity inhibitors (20 µM DNQX, 0.1 µM TTX, and 50 µM APV). The brains were digested with papain (Worthington Biochemical # LK003176, 50 units/ml, ∼2 µl per brain) at room temperature for 30 minutes. Brains were then resuspended and washed twice with ice-cold active SM medium after the digestion. To get the single cell suspension, we used flame-rounded 1000 µl pipette tips with different sized openings and triturated the brains until most of the tissues were dissociated. The resulting cell suspension was filtered by a 100 µm sieve. Hoechst dye (Invitrogen #R37605, one drop per 0.5 ml sample) was added into the sample tube in order to stain the nucleus before the single cell sorting. A BD Melody FACS machine in single-cell sorting mode was used for cell collection. Only the GFP and Hoechst positive single cells were collected. The collection devices were kept at 5 °C constantly during the sorting process.

### Single-cell RNA library preparation by modified CEL-Seq2

We used our previous modified CEL-Seq2 method for the single cell RNA sequencing of DANs around the clock. Briefly, single DANs were first sorted into 384 well plates prefilled with 0.6 µl primer mix (dNTP and primers), plates with sorted cells were centrifuged at 3000 *g* for 1 min at 4°C and then stored in −80°C until further processing. There are 96 ploy-T tailed primers in our method with which we can make 4 libraries from a 384 well plate. To increase the throughput, we used an Eppendorf EPmotion liquid handler to dispense first strand synthesis reagents and second strand synthesis reagents mixes. cDNA from the same primer set was pooled together, and cleaned by 0.8-fold AMPure beads before the in vitro transcription (overnight). Antisense RNA (aRNA) was converted to double stranded DNA with a random primer and T7-RA5 primer. The resulting cDNA underwent another final second round IVT step at 37°C overnight was followed by ExoSAP treatment (Affymetrix 78200) for 15 min at 37°C. Other steps were performed as described in the CEL-Seq2 protocol.

### Single-cell RNA library preparation by 10X Chromium

The same method was used to make the single cell suspension as described above. We first collected GFP and Hoechst positive single cells in a 1.5ml Eppendorf tube with 0.3ml collection buffer (PBS+ 0.04%BSA). The cells were spun down on a centrifuge by 700g for 10 minutes. We used the Chromium Single Cell 3′ kit (v3) of 10X Genomics. The libraries were prepared according to the standard User Guide (CG000315 Rev B) from 10X without any modifications.

### Library sequencing and raw data processing

Both the CEL-Seq2 libraries and 10X libraries were sequenced by Illumina Nextseq 550 with High Output Kit v2.5 (75 Cycles). zUMIs and Cellranger were used to map the sequencing data to the *Drosophila* genome (dm6) and count the reads from CEL-Seq2 and 10X Chromium separately(Parekh et al., 2018). Only the alignments to annotated exons were used for UMI quantitation.

The sequencing depth in CEL-Seq2 and 10X are different, so we used two different criteria to filter out low-quality cells in CEL-Seq2 and 10X experiments before the clustering analysis. For the cells in CEL-Seq2 experiment, we used the following criteria: (1) fewer than 500 or more than 6000 detected genes (where each gene had to have at least one UMI aligned); (2) fewer than 4000 or more than 75000 total UMI; (3) gene expression entropy smaller than 5.0, where entropy was defined as *-nUMI * ln(nUMI)* for genes with n*UMI >0*, where nUMI was a number of UMI in a cell. For the cells in 10X experiment, we used the following criteria: (1) fewer than 300 or more than 3500 detected genes (where each gene had to have at least one UMI aligned); (2) fewer than 1000 or more than 55000 total UMI; (3) gene expression entropy smaller than 5.0. We used Scrublet to detect the possible doublets in the 10X experiment, these cells were excluded from the following analysis.

### Dimensionality reduction and clustering

The method used for single cell clustering has been described previously (Ma et al., 2021). In short, we integrated the cells from different methods and time points using integration functions from the Seurat (version 3.0.2) package (Butler et al., 2018). First, we separated the single cell data by methods and time points and used the SCTransform function to transform data using the normalization and variance stabilization of counts. Batch effect was removed by regressing out numbers of genes, UMIs, detected genes per cell, sequencing batches, percentage of mitochondrial transcripts. We computed 3000 variable genes at each time point and method and found a subset of variable genes that were common to 8 conditions (6 time points from CEL-Seq2, 2 time points from 10X). From this set of common variable genes, we removed the mitochondrial, ribosomal, and transfer RNA genes. The resulting genes were used for integrating data using Seurat *FindIntegrationAnchors* and *IntegrateData* functions. Finally, we performed principal component analysis (PCA) on scaled gene expression vectors (z-scores) and reduced the data to the top 49 PCA components. This analysis resulted in 70 initial clusters, we next filtered the clusters based on the following criterion: first, all clusters must have cells from CEL-Seq2 and 10X, second, among the CEL-Seq2 data in each cluster, there should be cells from all time points throughout the day, third, clusters with low number of genes and transcripts were excluded. The cells in confident clusters were iterated one more time for the clustering as described above. We visualized the data using t-SNE embedding except where indicated specifically and reported a relative, normalized number of UMIs in a cell as TP10K – transcripts per 10 thousand transcripts.

### Differentially expressed genes in each cluster

The Seurat *FindAllMarkers* function with a negative binomial generalized linear model was used to identify the differentially expressed in each cluster. The p-values were adjusted for multiple hypothesis testing using Bonferroni method. We used an adjusted p-value significance of 0.05 and fold change cutoff of 1.25 as the threshold of significant differential expression.

### Matching single cell and bulk RNA sequencing in DANs

The bulk RNA sequencing results from different DAN subgroups were downloaded from (Aso et al., 2019). Only the results from FACS sorted samples were included in the current study. We first computed the enriched marker genes in DAN clusters by *FindAllMarkers* function from Seurat. The top 50 enriched genes from each cluster were used to compute the gene expression correlation between single cell and bulk RNA sequencing result.

### Immunohistochemistry

Immunohistochemistry was performed on 3–5 days old flies. Flies were fixed with 4% (vol/vol) paraformaldehyde with 0.5% Triton X-100 for 2 hours and 40 min at room temperature. Brains were dissected in 0.5% PBST and then washed twice (10 min) in 0.5% PBST buffer with rotation. 10% Normal Goat Serum (NGS; Jackson ImmunoResearch Lab) was used for blocking overnight at 4°C. Mouse anti-TH at 1:1000 dilution, rat anti-TIM at 1:200 dilution and chicken anti-GFP antibody at a 1:1000 were used as primary antibody and incubated with the brains overnight at 4°C, the brains were then washed twice (10 min) in 0.5% PBST buffer at room temperature. The corresponding secondary antibodies were added and incubated overnight at 4°C. Brains were mounted in Vectashield (Thermal Fisher) and imaged on a Leica SP5 confocal microscope. The images were processed by ImageJ.

### Data and code availability

The raw sequencing data and processed gene expression matrix have been deposited in GEO under accession number GSE198948. Custom codes for the analysis in the current study are available on GitHub: https://github.com/rosbashlab/scRNA_seq_Clk_vs_DANs.

## Acknowledgements

We thank the members of Rosbash lab for critical reading and thoughtful discussion of the manuscript. We greatly appreciate Drs. Liqun Luo, Larry Zipursky, Paul Garrity, Kai Zinn and Yerbol Kurmangaliyev for insightful comments on the manuscript.

---

**Supplementary Figure 1.**
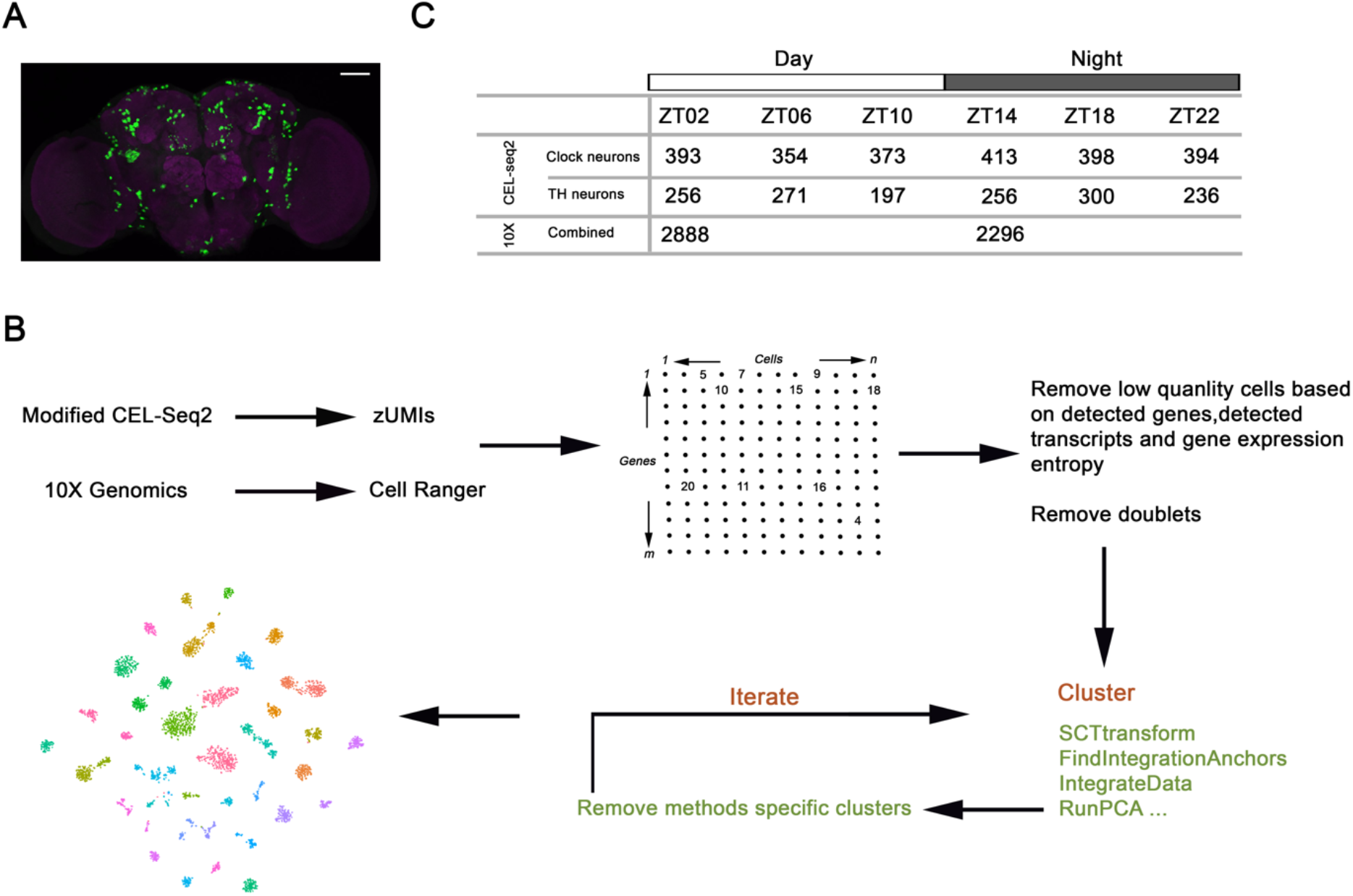
(A) Expression pattern of *Clk856-GAL4;TH-GAL4* > *UAS-Stinger-GFP* co-stained with anti-GFP (green) and nc82 (magenta). Scale bar is 50 μm. (B) Flow diagram showing the data processing. zUMIs and Cellranger were used to map and count single cell RNA libraries from CEL-Seq2 and 10X Chromium separately. The cells were filtered based on the number of detected genes, transcripts and gene expression entropy. Scrublet was used to identify possible doublets in 10X Chromium data; these doublets were excluded in the downstream analysis. (C) The number of high-quality cells from CEL-seq2 and 10X Chromium at each time point after initial filtering.

**Supplementary Figure 2.**
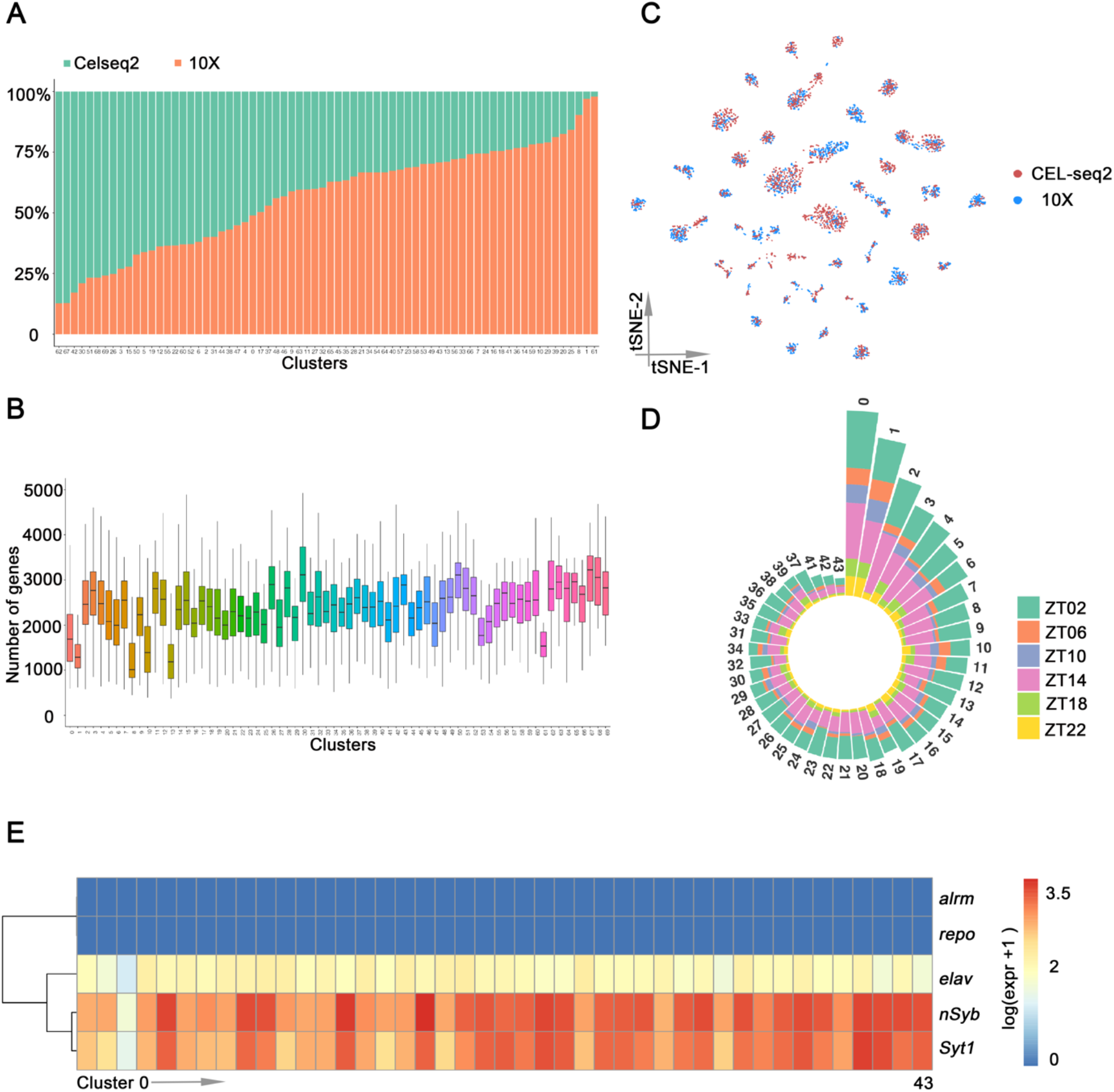
(A) The percentage of cells from CEL-seq2 and 10X Chromium in each cluster. The clusters are ordered by the percentage of cells from the CEL-Seq2 method. (B) Box plot showing the number of detected genes in each cluster. Each cluster has relatively similar numbers of genes with some exceptions. Numbers on the x-axis represent the 70 original clusters. (C) t-SNE plot showing the cells from CEL-seq2 (red) and 10X Chromium (blue) in the final 43 high confidence clusters. (D) Circled bar plot showing that in high confidence clusters there are cells from 6 time points in Light: Dark conditions. (E) Heatmap showing the glial and neuronal marker gene expression in all clusters.

**Supplementary Figure 3.**
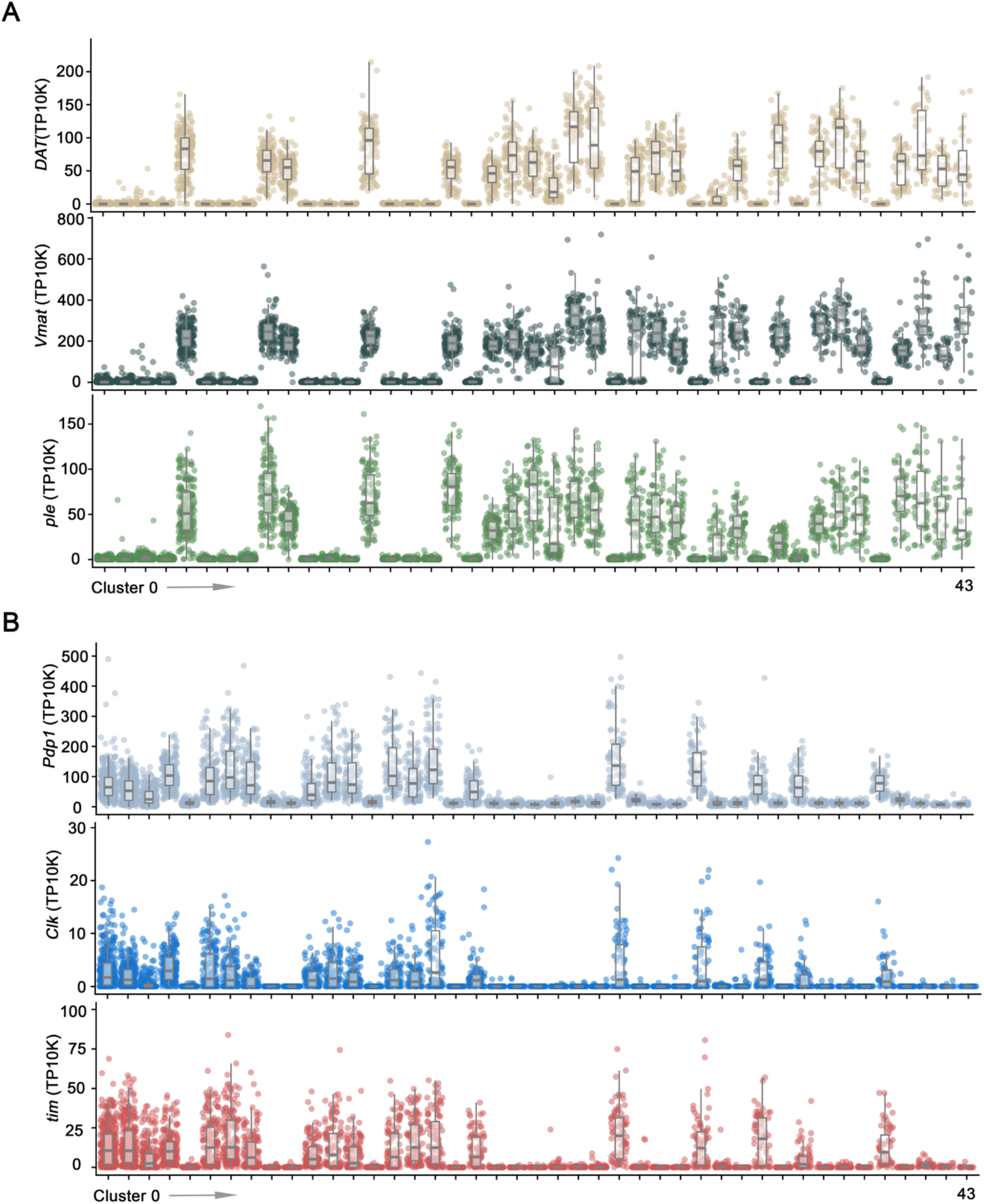
(A-B) Dot plot showing marker gene expression in DAN (A) and clock neurons (B) clusters. *ple*, *Vmat* and *DAT* are enriched in DANs. *tim*, *Clk* and *Pdp1* are exclusively expressed in clock neurons. Gene expression levels for each cell were normalized by total expression level; we report transcripts per 10 thousand transcripts (TP10K). Clusters are ordered by size.

**Supplementary Figure 4.**
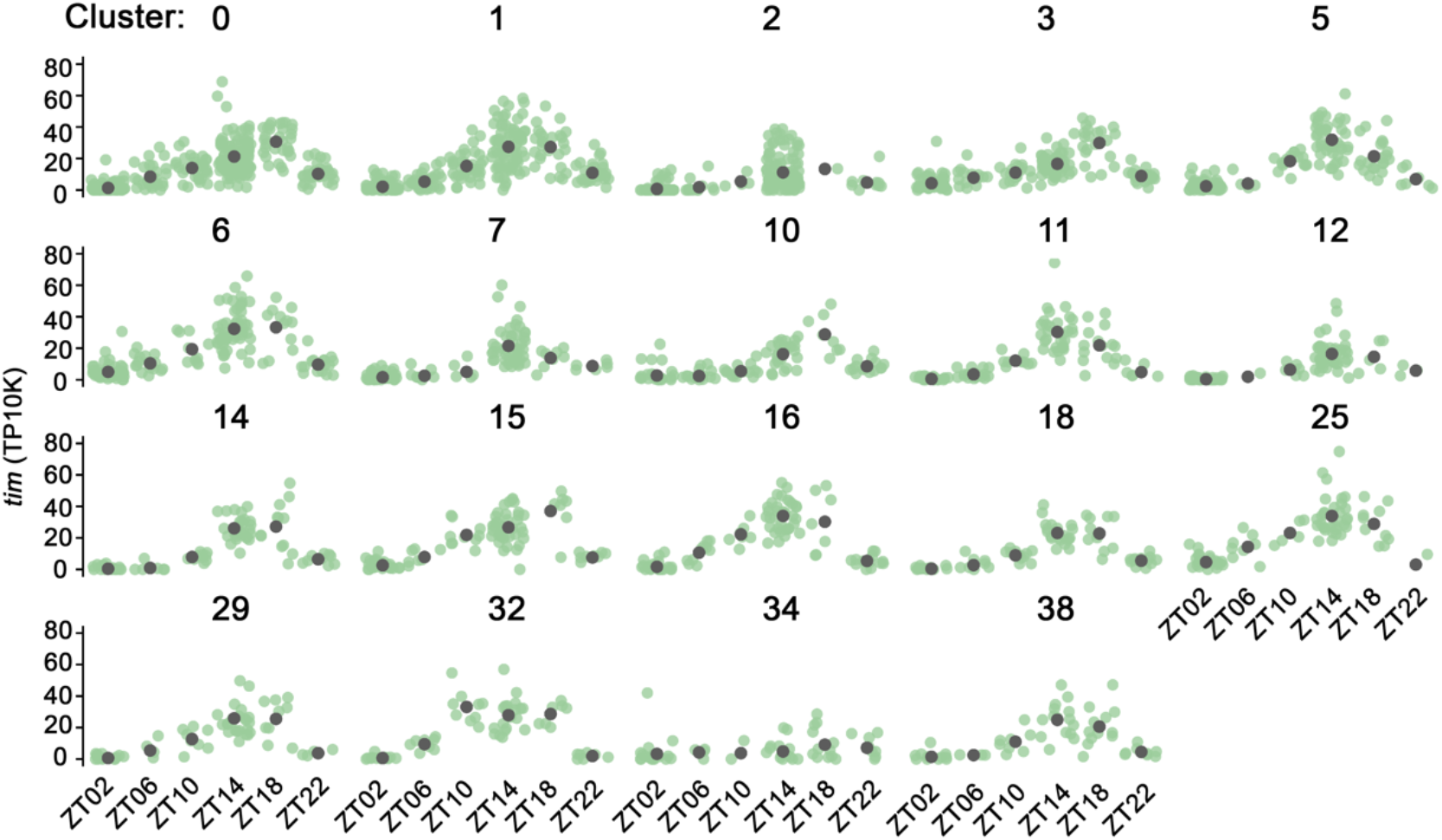
*tim* expression in all clock neuron clusters throughout the day in Light: Dark cycles. The green dots represent single cells and the gray dots represent the average *tim* expression in each cluster. Gene expression levels for each cell were normalized by total expression level.

**Supplementary Figure 5.**
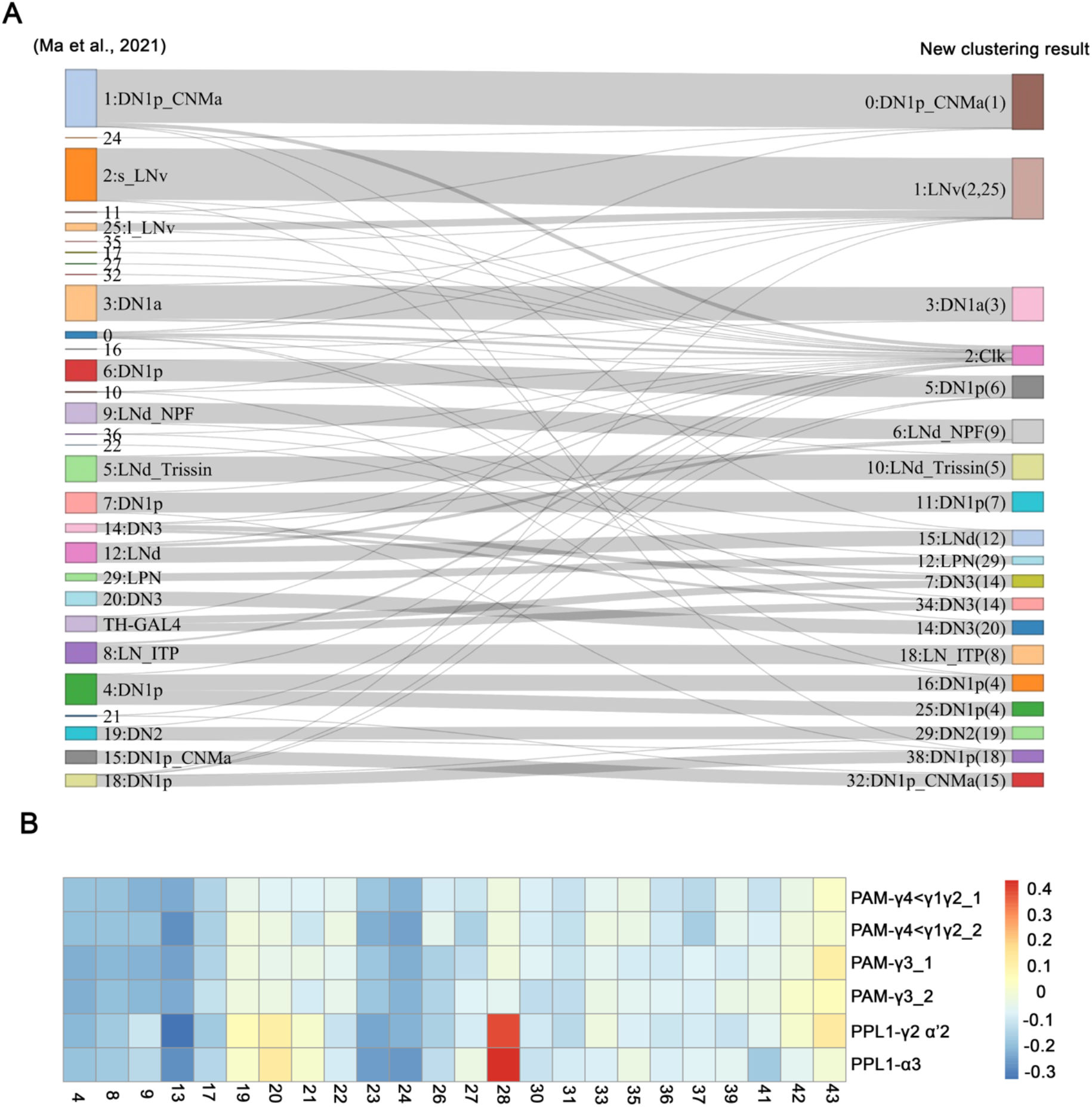
(A) Sanky plot showing the contribution of predefined clock neurons clusters (left) to the classification (right) in the current study. Each node represents a single cell cluster. For comparison, each clock neuron cluster retains its original identifying number in the parentheses as it was reported previously. (B) Heatmap showing the gene expression correlation between single cell clusters and different DAN subgroups. Only the transcriptomic results from FACS sorted cells were included in the analysis.

**Supplementary Figure 6.**
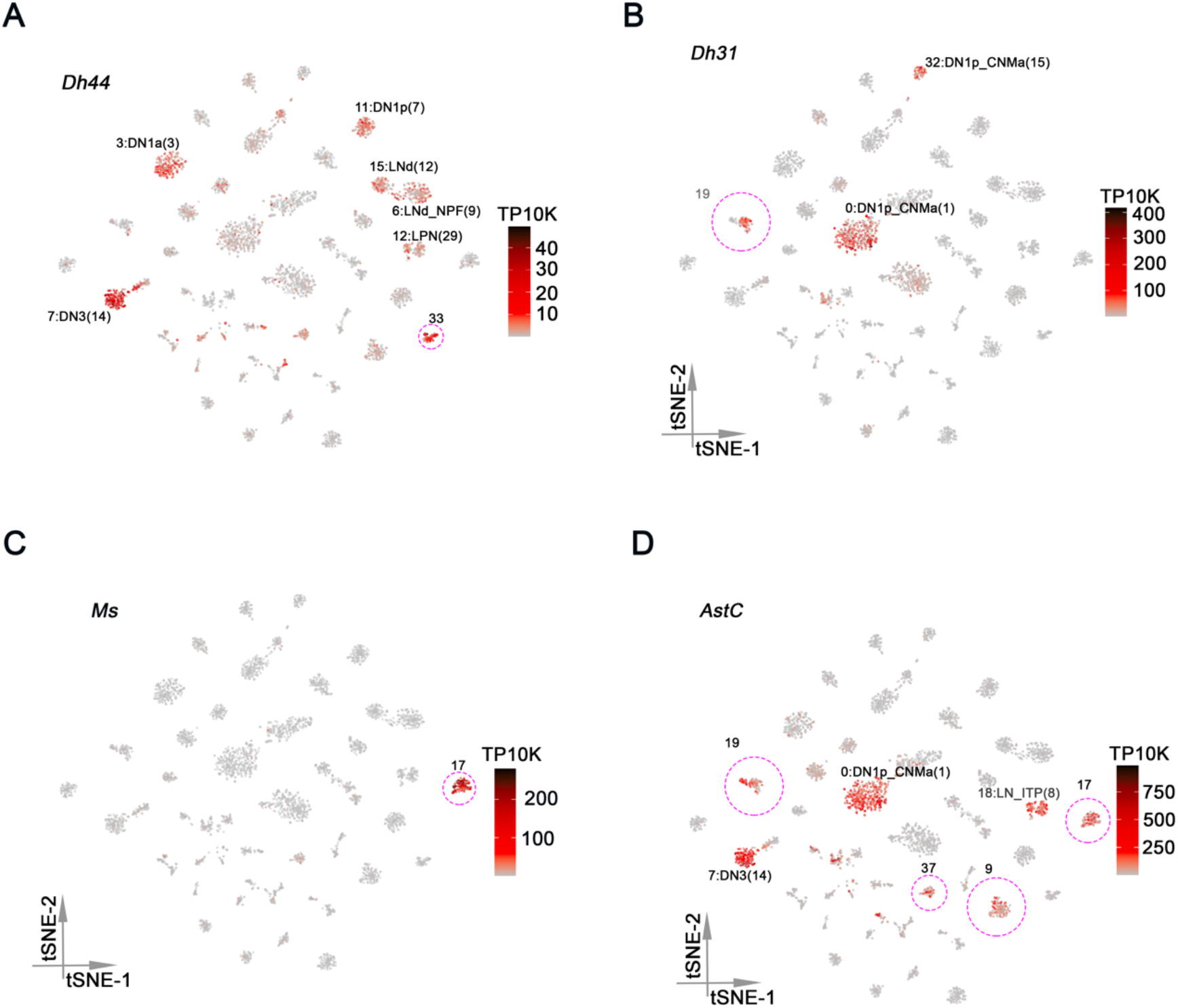
(A-D) t-SNE plots showing *Dh44* (A), *Dh31* (B)*, Ms* (C) and *AstC* (D) expression in all clusters. Each cell is colored by the expression level with red indicating high expression and gray indicating low expression. The DAN clusters are highlighted by dashed red circles.

**Supplementary Figure 7.**
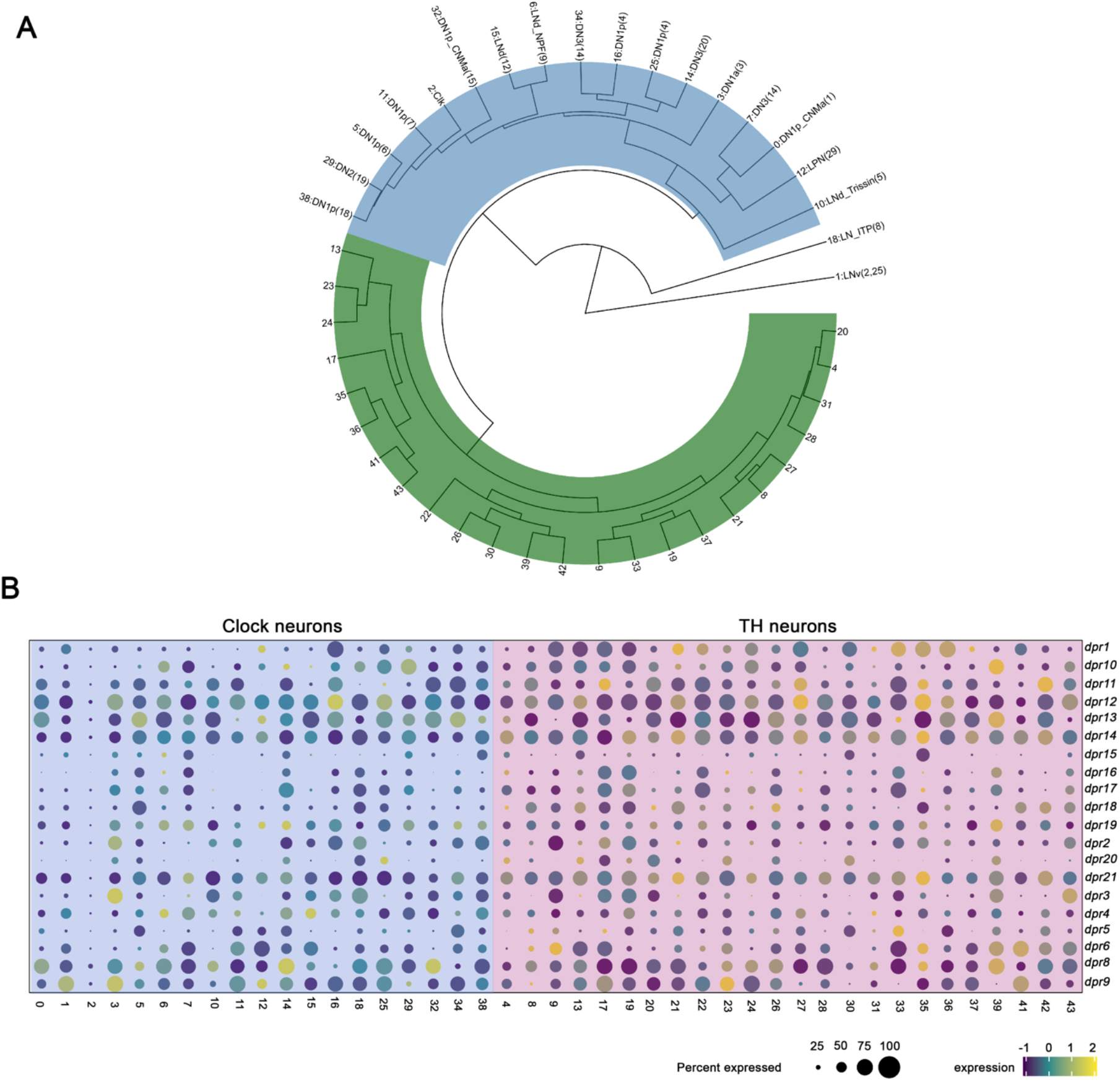
(A) Hierarchical clustering of identified clusters based on 338 highly variable genes. The analysis was based on integrated dataset. (B) Dot plot showing the *Dpr* family gene expression in clock neuron (left) and DANs (right) clusters. Size of dot indicates what percentage of cells in a particular cluster that express the indicated DIP-family member. Color indicates the mean expression within that cluster.

**Supplementary Figure 8.**
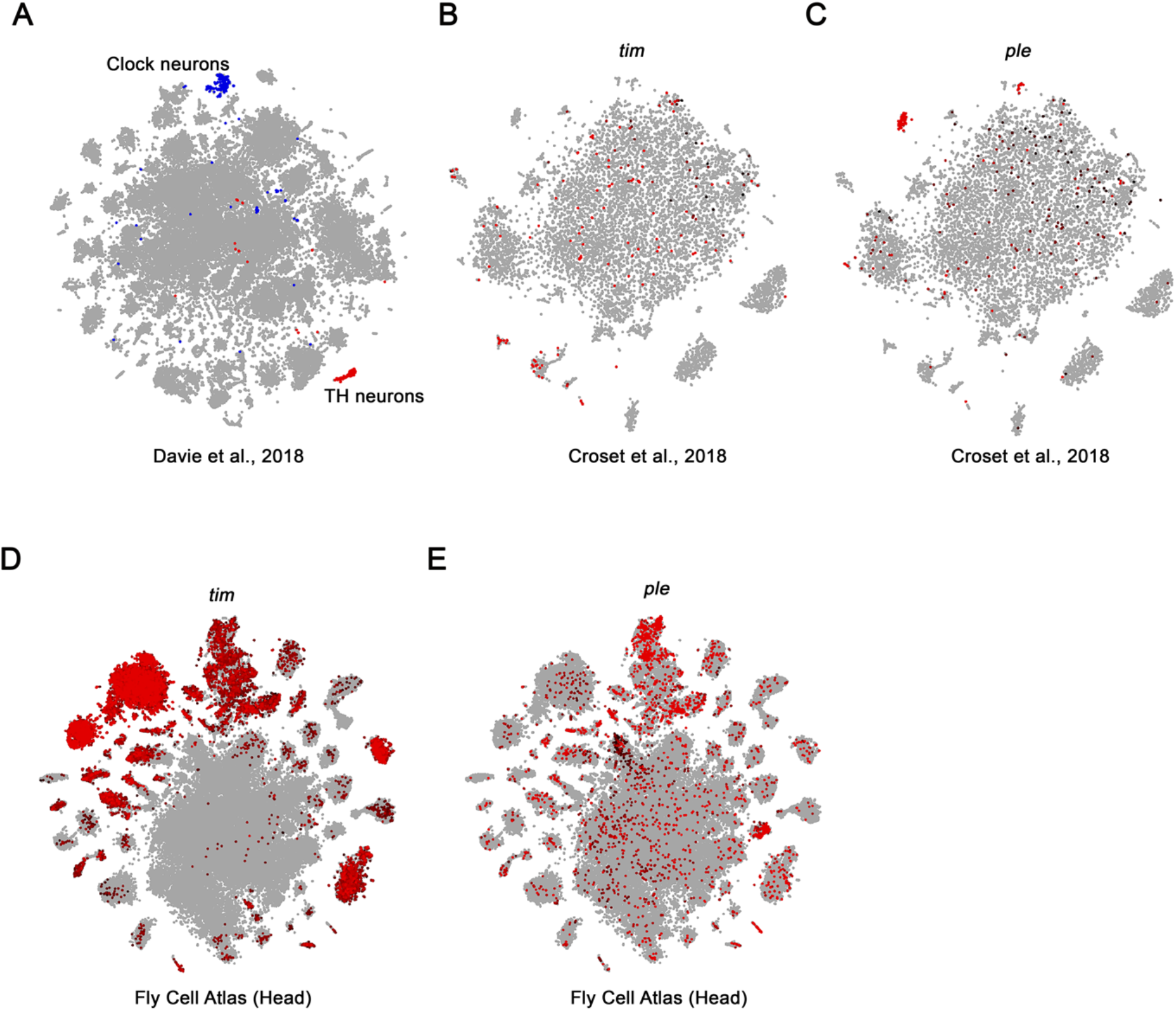
(A) t-SNE plots showing the identified clock neuron (blue) and DAN (red) clusters from (Davie et al., 2018). (B-C) *tim* (B) and *ple* (C) expression in (Croset et al., 2018). (D-E) *tim* (D) and *ple* (E) expression in Fly Cell Atlas head result (Li et al., 2022).

